# *In silico* analysis of the invasion mechanics and invasiveness of the plasmodium falciparum merozoite

**DOI:** 10.1101/2025.06.26.661885

**Authors:** Chimwemwe Msosa, Tamer Abdalrahman, Thomas Franz

## Abstract

Although there has been considerable progress in understanding the factors that determine the invasiveness of plasmodium falciparum merozoites, the collective role of the biophysical characteristics of erythrocyte deformability in the invasion process is poorly understood. Cell shape, cytoplasmic viscosity, and membrane stability are the main determinants of erythrocyte deformability, but it remains unknown how these properties affect the merozoite invasiveness. This study aimed to investigate computationally (i) the role of erythrocyte morphology and merozoite-induced erythrocyte membrane damage in merozoite invasion and (ii) the suitability of mechanical markers of merozoite-induced erythrocyte membrane damage for screening of invasion-blocking antimalarial drugs. Finite element models were developed to represent a human erythrocyte and a spherocyte, their invasion by a malaria merozoite, and erythrocyte compression and nanoindentation as mechanical assays for membrane damage. Smoothed particle hydrodynamics represented the erythrocyte cytoplasm, and merozoite-induced erythrocyte membrane damage was implemented with a constitutive model. The invasiveness of the merozoite decreases with increased erythrocyte sphericity associated with genetic disorders such as hereditary spherocytosis. The invasiveness is larger when membrane damage is induced in the erythrocyte at an early invasion stage than throughout the invasion process. The minimum force required for a malaria merozoite to invade a human erythrocyte was predicted to be 11 pN. The findings on the invasion mechanics can guide future studies into the invasiveness of the merozoite. The nanoindentation simulations point to the potential of nanoindentation to determine erythrocyte membrane damage for screening novel invasion-blocking anti-malaria drugs.

## 1. Introduction

The invasion of malaria merozoites into human erythrocytes has been extensively studied as a potential target for antimalarial drugs (Flannery et al. 2013). During the invasion of erythrocytes, the merozoite is highly exposed to the host immune system and is highly vulnerable to therapeutic interventions; hence, it is an essential target for antimalarial drugs. Despite the considerable interest in studying the invasion process, fundamental gaps remain in our understanding of the invasion process. Despite decades of research, an incomplete understanding of the disease’s physiopathology continues to hinder eradication efforts. Beyond oxidative stress-mediated damage (Percário et al. 2012), erythrocyte invasion by the Plasmodium falciparum merozoite involves highly coordinated ligand–receptor interactions that induce erythrocyte membrane remodelling. The binding of parasite ligands such as EBA-175 to glycophorin A activates intracellular phosphorylation cascades involving cytoskeletal proteins, leading to altered mechanical properties of the erythrocyte membrane (Sisquella et al. 2017). Pharmacological inhibition of this remodelling pathway has been shown to suppress phosphorylation cascades, limit changes in membrane deformability, and block merozoite invasion, emphasising the importance of host membrane mechanics in conditioning erythrocytes for successful parasite entry (Sisquella et al. 2017). To date, the impact of erythrocyte morphology and local erythrocyte membrane damage on the invasiveness of merozoites has not been comprehensively explored due to the lack of appropriate and detailed invasion mechanics models. As such, *in silico* invasion mechanics models offer an alternative approach for investigating the invasiveness of the merozoite, complementing *in vitro* and *in vivo* studies.

Various *in silico* models have been developed to describe erythrocyte mechanics. These include cortical shell Newtonian liquid drop models applied to biological cells such as erythrocytes(Lim et al. 2006), finite element models of erythrocyte deformation under shear stress (Ahmad and Ahmad 2015), and microstructural representations of the erythrocyte spectrin network (Fai et al. 2017). Multiscale approaches based on dissipative particle dynamics have further enabled simulation of infected erythrocyte deformation across intra-erythrocytic developmental stages by coupling subcellular and vascular-scale mechanics. More recently, atomistically enriched constitutive models combined with mesh-free numerical methods have demonstrated improved predictive accuracy of erythrocyte deformability (Ademiloye et al. 2018). However, despite these advances, none of these models have been applied to investigate the invasion mechanics of malaria parasites partly due to limited validation data for mechanical and other multiscale models.

The merozoite entry into an erythrocyte is an active process that involves the application of actomyosin-based forces on the erythrocyte membrane. The forces are transmitted to the erythrocyte membrane through contact with the merozoite surface (Dasgupta et al. 2014). To date, a detailed analysis of the mechanistic role of the erythrocyte membrane and associated structure, i.e., the spectrin network involved in the invasion process, is limited to 2D analytical models (Abdalrahman and Franz 2017). Additionally, current analytical models do not incorporate large deformations and remodelling of the erythrocyte membrane, which limits their application to early-stage invasion with a maximum invagination depth of 10% of the merozoite length. Thus, there is a need to develop realistic 3D invasion models that account for all factors that determine the merozoite invasiveness, i.e., merozoite-induced membrane damage.

Hereditary spherocytosis is caused by genetic alteration of erythrocyte membrane proteins, leading to the formation of spherocytes. Previously, it has been documented that cells with hereditary spherocytosis have abnormal protein structure and thus have a low susceptibility to infection by the merozoite (Eber and Lux 2004). Despite this finding, little is known about the higher invasion resistance of these cells.

The current study aimed to computationally investigate the erythrocyte mechanics during malaria parasite invasion with emphasis on (i) the factors contributing to the merozoite invasiveness and (ii) the impact of local disruption of the spectrin network on the global mechanical properties of the erythrocyte for assessing the feasibility of mechanical markers for testing the efficacy of invasion-blocking antimalarial drugs.

## 2. Materials and methods

### 2.1. Geometric modelling

#### 2.1.1. Healthy erythrocyte

The initial biconcave geometry of the erythrocyte was defined by the following equation (Evans and Fung 1972):

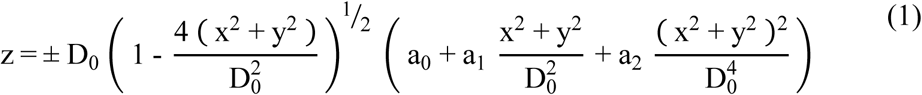

with principal coordinate directions x, y, z, the diameter of the undeformed erythrocyte D_0_ = 7.82 µm, and shape parameters a_0_ = 0.0518, a_1_ = 2.026 and a_2_ = –4.491 (Figure 1 a). The generated erythrocyte model has a volume of 94.47 µm^3^ and a surface area of 135 µm^2^ consistent with the literature (Li et al. 2014).

**Figure 1:**
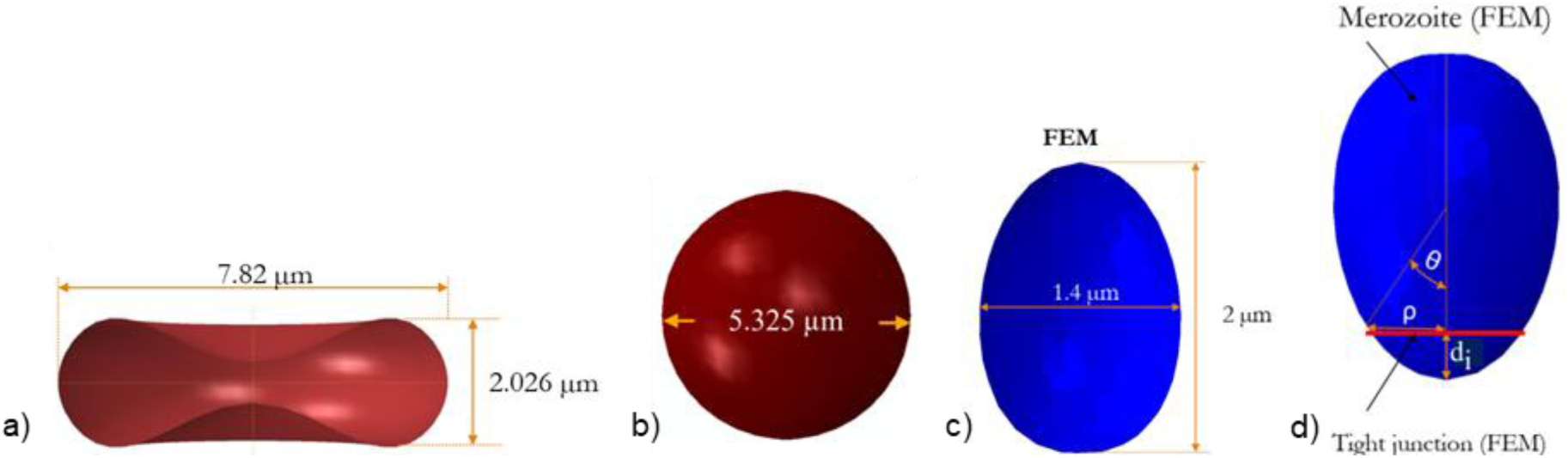
a) Erythrocyte geometry, b) Spherocyte geometry, c) Dimension of Plasmodium Falciparum merozoite based on cryo-EM data from Dasgupta et al. (2014, Fig. 2), d) Geometry of rigid egg-shaped merozoite used in current study.

#### 2.1.2. Spherocyte

In individuals with spherocytosis, erythrocytes take a spherical form due to alterations of erythrocyte membrane proteins. The geometry was described with

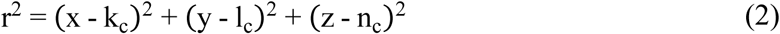

where r = 2.66 µm is the radius of the spherocyte (Li et al. 2016) and k_c_, l_c_, n_c_ are centre coordinates of the spherocyte, where k_c_ = l_c_ = n_c_ = 0. x, y, and z denote the coordinate points on the surface of the spherocyte (Figure 1 b). The surface area to volume ratio is 14% smaller for the spherical shape than the discoid geometry.

#### 2.1.3. Plasmodium falciparum merozoite

The merozoite shape has been previously described based on cryo-x-ray images of free merozoites (Dasgupta et al. 2014, Fig. 2). From these data, the mean physical dimensions of the merozoite were determined as follows: Length L_m_ = 1.98 ± 0.08 μm, width W = 1.40 ± 0.06 μm, volume V_actual_ = 1.71 ± 0.15 μm^3^ and surface area A_actual_ = 8.06 ± 0.72 μm^2^ (Figure ^1^ c). The 3D merozoite geometry (Figure 1 d) was generated with

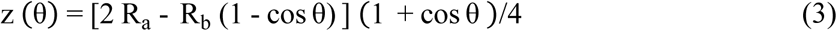

**Figure 2:**
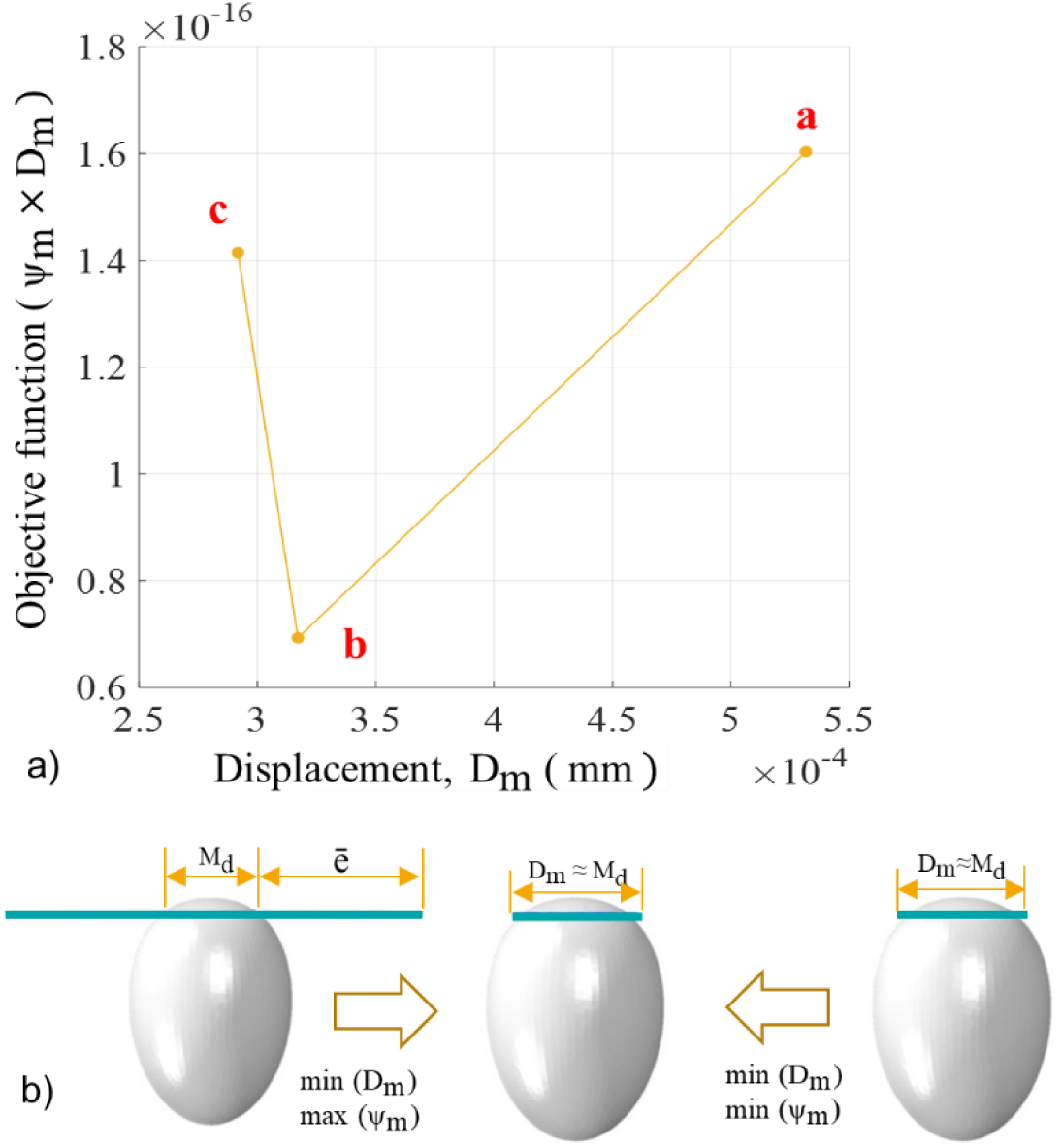
Illustration of a) the minimisation process of the objective function and b) the determination of the Mooney Rivlin parameters.

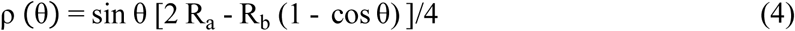

where ρ and z are Cartesian coordinates, R_a_ = 1 µm, R_b_ = 0.7 µm are parameters that determine the egg shape profile of the merozoite, and θ is a polar angle with 0° ≤ θ ≤ 180°.

Images from electron microscopy, cryo-electron tomography, cryo-x-ray tomography and widefield deconvolution fluorescence imaging of a merozoite during invasion show negligible changes in merozoite shape throughout the invasion process (Zuccala et al. 2016). Hence, this study treated the merozoite geometry as a rigid body.

#### 2.1.4. Tight junction between the merozoite and erythrocyte membrane

The merozoite pulls itself into the erythrocyte through the tight junction complexes, which it establishes after forming the invasion pit (Riglar et al. 2011; Preiser et al. 2000; Pinder et al. 2000). The tight junction was modelled as a deforming mechanical link between the merozoite surface and the erythrocyte membrane, forming an annulus-like structure to facilitate erythrocyte membrane wrapping. The annulus is defined as a circular ring with an internal diameter of 0.76 µm and a cross-sectional radius of 0.04 µm (Figure 1 d).

### 2.2. Constitutive modelling

#### 2.2.1. Erythrocyte membrane with merozoite-induced damage

During merozoite invasion, the erythrocyte membrane deformation was considered as mainly due to the mechanical loads exerted by the merozoite’s actomyosin machinery and other external sources, such as blood pressure, whereas the entropic deformation was considered negligible. Hence the Helmholtz free energy function for the erythrocyte membrane deformation was only represented as internal strain energy.

The damage induced by the merozoite was modelled by modifying the strain energy density function of the erythrocyte membrane. Since the erythrocyte membrane comprises primarily an elastic spectrin network and can be considered an elastic, isotropic, and nearly incompressible continuum, the strain energy density function is usually presented in a decoupled form comprising deviatoric and isochoric terms (Li 2016).

The combination of incompressibility and large deformation of a nearly incompressible hyperelastic material presents difficulties for a displacement-based finite element method as the constraint J = det F = 1 on the deformation field is highly nonlinear (Weiss 1994). To overcome this challenge, a displacement-based finite element scheme must invoke a small change measure of volumetric deformation. Consequentially, the deformation gradient was decomposed into the dilatational and deviatoric parts to apply separate numerical treatments to each part (Weiss 1994).

Therefore, the deformation gradient **F** (Gilson and Crabb 2009)and the left Cauchy-Green strain tensor **B** were divided into the volume-changing (dilatational) and the volume-preserving (distortional) parts, an approach often used in elasto-plasticity (Ogden 1978). The strain energy density function of the isotropic erythrocyte membrane was expressed in terms of the left Cauchy deformation tensor as

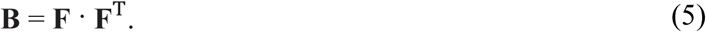

With **F** = **R U** and **F**^T^ = **R**^T^ **U**^T^, where **R** is a rotation matrix, and **U** is a stretch tensor, Eqn. (5) becomes

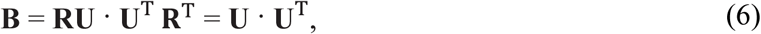

showing that the left Cauchy deformation tensor **B** is a stretch tensor and isotropic.

Hence the strain energy density function ψ of a damaged erythrocyte membrane can be written in terms of invariants of the left Cauchy-Green deformation tensor:

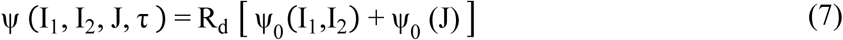

with

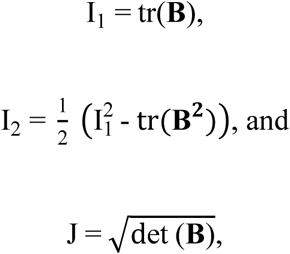

where **B** is the left Cauchy-Green tensor, R_d_ is the damage parameter, ψ_0_ is the strain energy density of an intact erythrocyte membrane, I_1_ and I_2_ are the first and second deviatoric strain invariants, respectively, while J represents the determinant of the deformation gradient and quantifies the local volumetric change, and τ is the indentation time in seconds.

A continuum damage mechanics framework for material stiffness deterioration suitable for implementation in Abaqus Explicit was used to simulate chemical damage induced by a merozoite during the entry process. The model draws on concepts from various strain-based damage models for soft biological materials and biodegradable polymers, which describe constitutive hydrolytic degradation and time-dependent behaviour (Vieira et al. 2011).

R_d_ is a function of chemical and mechanical damage parameters defined as:

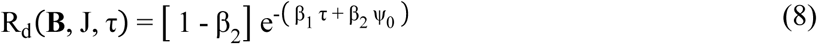

where **B** and J denote the left Cauchy green tensor and the volumetric strain of the intact erythrocyte membrane, respectively, τ is the indentation time, and ψ_0_ is the strain energy density per reference volume of the intact erythrocyte membrane, β_1_ is the chemical damage parameter, and β_2_ is the mechanical damage parameter.

For β_2_ = 0, the damage mode is purely chemical, i.e., damage to the erythrocyte membrane is not due to deformation. Thus, β_1_ represents chemical damage due to various amounts of phosphorylation, and β_2_ represents mechanical damage associated with the tearing or rupturing of the protein chains in the erythrocyte membrane skeleton. The erythrocyte’s membrane near incompressibility was defined with a Poisson’s ratio ν = 0.499. For incompressible materials, the contribution of the volumetric strain energy density function is neglected since J = 1. Since the erythrocyte membrane is nearly incompressible, the deviatoric strain invariants are

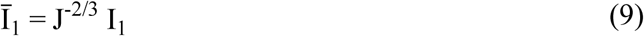

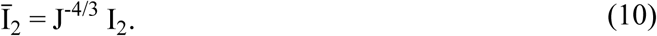

The variation of the strain energy potential δW_i_ is, by definition, equal to the internal virtual work per reference volume V_0,_ and can be written as (Dassault Systèmes Simulia Corp 2015):

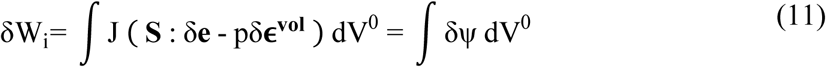

where

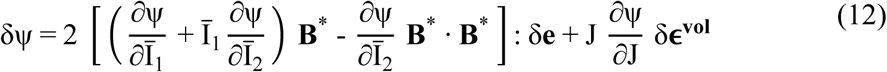

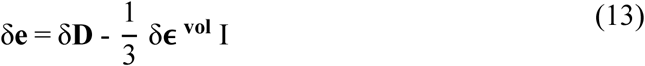

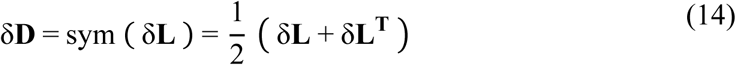

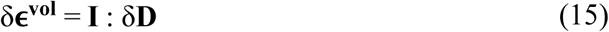

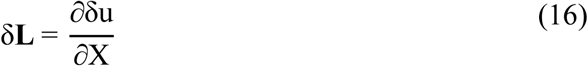

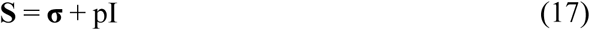

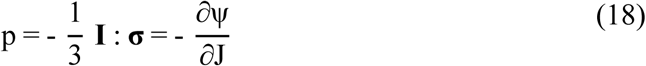

Here, **B\*** = J^-2/3^**B** is the corrected left Cauchy Green tensor, S is the deviatoric strain tensor, δe is the virtual deviatoric strain tensor, δε^vol^ is the virtual volumetric strain tensor, p is the hydrostatic pressure, δD is the virtual strain rate tensor, and δL is the virtual velocity gradient tensor.

Hence, the deviatoric stress with damage can be rewritten as:

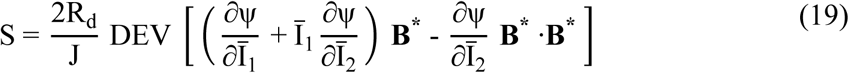

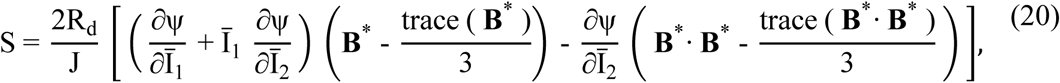

Therefore, the stress (deviatoric stress and volumetric stress) with damage can be written as

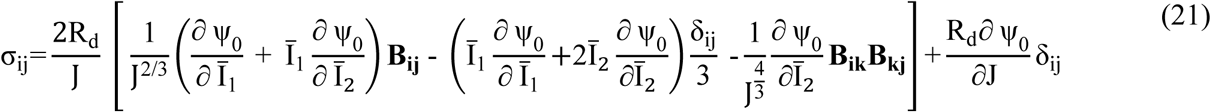

and further as

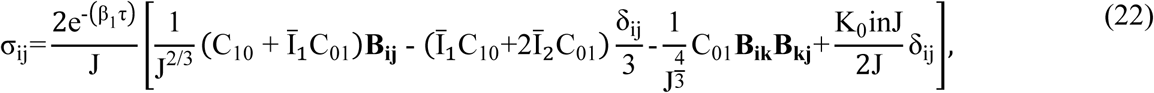

where δ is the Kronecker delta function, and ψ_0_ is the strain energy density function per unit reference volume of the intact erythrocyte membrane.

The developed erythrocyte membrane damage (EMD) model, Eqn. (22), to induce localised damage in the erythrocyte membrane was implemented using the VUMAT subroutine, whereas the Mooney Rivlin law model to describe the constitutive response of the intact erythrocyte membrane was implemented using the Abaqus materials module.

The erythrocyte membrane damage model was verified using a single shell element model subjected to an equi-biaxial strain of 1.1. The verification involved comparing the true stress obtained with the VUMAT subroutine and the built-in Mooney Rivlin law for an intact erythrocyte membrane with the chemical damage parameter β_1_ = 0. For this case, the constitutive responses of the developed VUMAT subroutine and the built-in Mooney Rivlin law are expected to be identical. Thereafter, the single-shell-element model was used with various degrees of chemical damage to evaluate the stability of the developed erythrocyte membrane damage model using Drucker’s stability criterion.

The material’s relative compressibility also determines the mechanical response of the erythrocyte membrane. The relative compressibility is the ratio of the initial bulk modulus K_0_ to the initial shear modulus µ_0_ of the material:

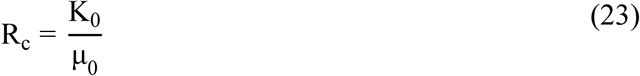

where µ_0_ and K_0_ are defined by Eqns. (24) and (25), respectively, and large R_c_ values show that the material is less compressible:

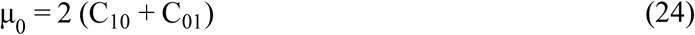

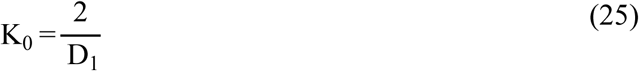

The Poisson’s ratio ν for hyperelastic materials is related to R_c_ by:

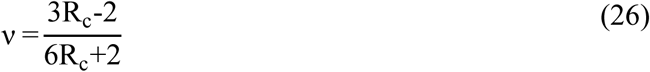

The inverse bulk modulus D_1_ defines the material compressibility and is expressed as:

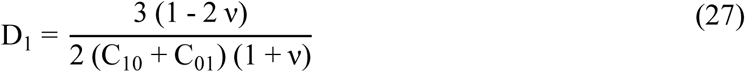

For this study, the Poisson’s ratio of the erythrocyte membrane was set to 0.499 to avoid numerical singularity; hence, D_1_ was set to 12 mm^2^/N. The material parameters for the Mooney Rivlin model are computed from the elastic modulus by:

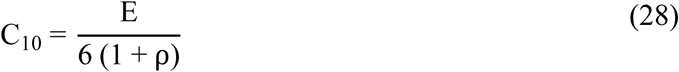

and C_01_ = ρ C_10_ (Zhang and Zhang 2011). For an elastic modulus E = 1 kPa and ρ = 0.1 used in the current study, the corresponding values of C_10_ and C_01_ are 152 Pa and 15.2 Pa, respectively.

#### 2.2.2. Tight junction

The tight junction was represented as an annulus, mimicking its function during the invasion process. The energy associated with the work done by the tight junction is not yet known. However, an estimate of the minimum energy contribution of the tight junction required for a successful invasion was determined with the developed finite element model, and the Mooney Rivlin law was used to define the mechanical response of the annulus structure. The material parameters were determined by minimising the objective function defined by:

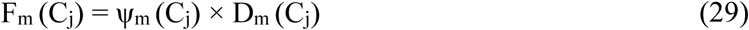

where ψ_m_ is the strain energy density function for the annulus structure, and D_m_ is the diameter of the tight junction at maximum indentation depth at the end of the invasion when the simulation step time is 1.1 s. The binary search algorithm below was used to search for material parameters that minimise the objective function in Eqn. (29).

Given an array C_j_ of n elements C_0j_, C_1j_, C_2j_ …. C_nj-1_ such that C_0j_ ≤ C_1j_ ≤ C_2j_ ≤ … ≤ C_nj-1,_ the following pseudo-code uses the binary to find the value of the material parameter in C_j_ that minimises the objective function.

1. Set L_j_ = 0 and R_j_ = n_j_ – 1
2. If L_j_ > R_j_, the search terminates as unsuccessful
3. Determine the middle element index m_j_ = floor ([L_j_ + R_j_]/2)
4. Compute the D_m_ using the middle element parameter in C_j_.
5. If D_m_ > M_d_ + ē update C_j_ such that C_j_ ≥ C_m_, compute F_m_ and go to step 2.
6. If D_m_ < M_d_ + ē, update C_j_ such that C_j_ ≤ C_m_, compute F_m_ and go to step 2.
7. If D_m_ ≈ M_d_ the search is done, and compute F_min_ = min (F_m_)

Here, M_d_ is the width of the merozoite at maximum indentation depth, and ē is the clearance between the tight junction and the merozoite at the maximum indentation depth. When F_m_ = F_min_ = min (F_m_) (Figure 2 a), C_j_ gives a minimum strain energy ψ_m_ (C_j_) such that D_m_ ≈ M_d_ (Figure 2 b). The strain energy ψ_m_ (C_j_) of the annulus structure increases from point a to b and c (Figure 2 a) while the diameter D_m_ decreases from point a to b but remains constant from point b to c (Figure 2 a and b).

The mechanical properties of the annulus structure that mimics the tight junction were determined by minimising the objective function, yielding Mooney-Rivlin parameters C10 = 0.04 MPa, C01 = 0.004 MPa, and D_1_ = 0.3 mm^2^/N. The determined mechanical properties represent the minimum energy required by the annulus structure to ensure erythrocyte membrane wrapping.

#### 2.2.3. Erythrocyte cytoplasm

The smoothed particle hydrodynamics (SPH) method was used to model the erythrocyte cytoplasm deformation. Smoothed particle hydrodynamics is a fully Lagrangian mesh-free modelling scheme permitting the discretisation of a prescribed set of continuum equations by interpolating the properties directly at a discrete set of points distributed over the solution domain. This approach was first developed to solve PDE problems in astrophysics (Gingold and Monaghan 1977). In Abaqus, the SPH scheme discretises the continuum partial differential equations (Violeau and Rogers 2016). SPH uses an evolving interpolation scheme to approximate a field variable at any point in a domain. Using the particle approximation or field function Eqn. (30) and its derivative Eqn. (31), the Navier-Stokes equation is discretised and solved using the explicit time integration method.

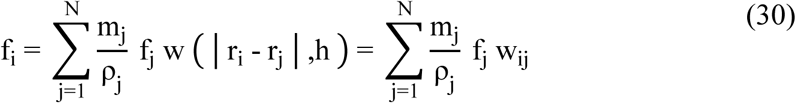

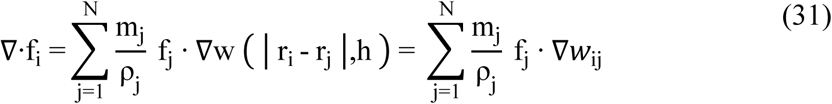

where N is the total number of particles, h is the smoothing length, and r_i_ and r_j_ are the position vectors of the particle of interest and the particle in the neighbouring region, respectively. The field function f and its derivative are constructed using a smoothing or kernel function w (Wang et al. 2016, Fig. 1). Thus, the value of a variable at a particle of interest can be approximated by summing the contributions from a set of neighbouring particles, denoted by subscript j, for which the “kernel” function, w, is not zero.

In Abaqus, the erythrocyte cytoplasmic domain was converted to SPH particles by activating the built-in conversion functionality. The erythrocyte cytoplasm primarily comprises viscous haemoglobin, mathematically described by the Navier-Stokes equation in the Lagrangian form (Ye et al. 2015). The erythrocyte cytoplasm is generally considered an incompressible Newtonian liquid, and thus, its dynamics are predicted by using the Navier-Stokes equations given by:

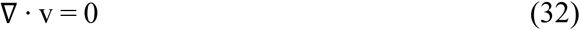

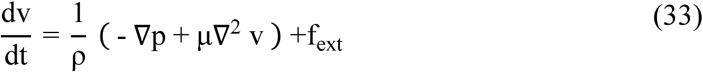

where p, v, ρ, µ and f_ext_ represent the pressure, velocity, density, dynamic viscosity and external force vector, respectively. SPH solves the Navier-Stokes equations by discretising the whole computational domain into a set of particles. The Mie-Grüneisen equation of state, Eqn. (34), was used to model incompressible viscous laminar flow governed by the Navier-Stokes equation of motion. The volumetric response is governed by the equations of state, where the bulk modulus acts as a penalty parameter for the incompressible constraint. Since the viscosity of the erythrocyte cytoplasm is small, a small amount of shear resistance was specified in the materials module to suppress shear modes that can otherwise tangle the mesh. Here, the shear stiffness or viscosity was used as a penalty parameter. The default hourglass control was used because when the shear model is defined, the hourglass control forces are calculated based on the shear resistance of the erythrocyte cytoplasm, which provides very low shear strength, insufficient to prevent spurious hourglass modes. An equation of state is necessary for the erythrocyte cytoplasmic domain to link pressure P and density ρ. The Mie-Grüneisen equation of state used for this purpose (Monaghan 1988) is given as follows:

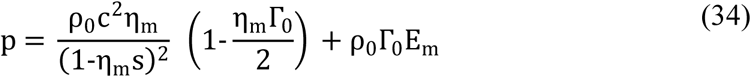

where ρ_0_ is the reference density, and c is the speed of sound, Γ_0_ = 0 is a material parameter, η_m_ = 1 − ρ_0_/ρ is the nominal volumetric compressive strain, E_m_ is internal energy per unit mass. The background pressure P_0_ is added to avoid negative pressure values. The density is estimated from the particle distribution utilising the SPH interpolation. c and s define the linear relationship between the shock wave velocity, U_s_, and the particle velocity, U_p_, as follows:

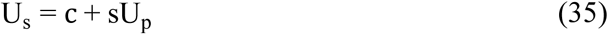

where s was set to zero such that U_s_ = c = 1,000 mm/s.

### 2.3. Finite element meshes

The finite element model comprised multiple components with varying element lengths: the erythrocyte membrane, the tight junction, the rigid merozoite, and the cytoplasm (Figure 3 a-d).

**Figure 3:**
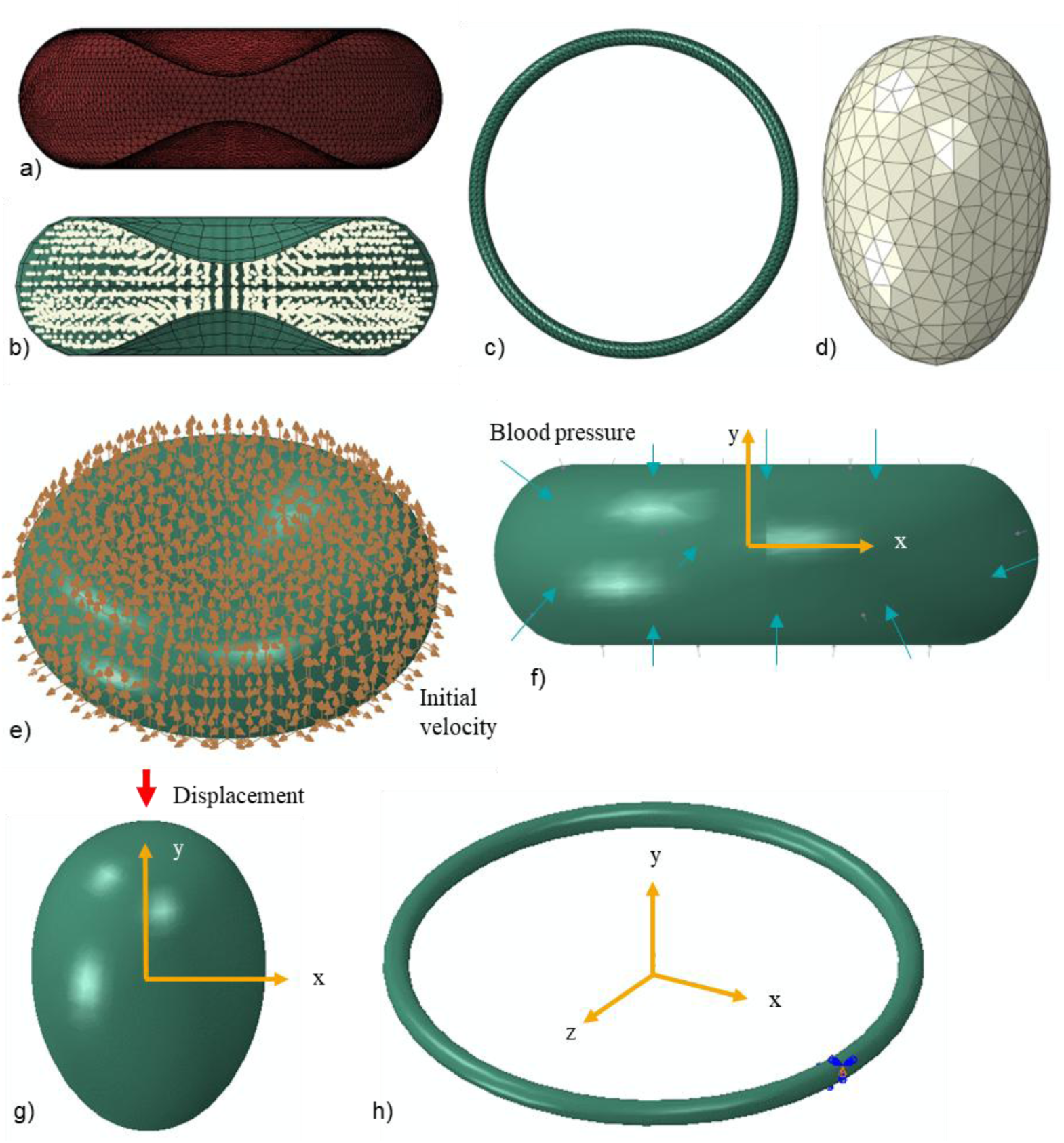
Finite element meshes and boundary conditions. Finite element mesh of a) erythrocyte membrane with triangular shell elements, b) erythrocyte cytoplasm with 8-node linear brick elements with reduced integration and hourglass control (C3D8R), c) annulus structure of the tight junction with ten-node modified quadratic tetrahedron elements (C3D10M), and d) rigid merozoite with three-node 3D rigid triangular elements (R3D3). Boundary conditions: e) Blood pressure applied on the outer surface of the erythrocyte membrane, f) the initial velocity of 2×10^-30^ mm/s applied at each node of the erythrocyte membrane, g) each node of the annulus structure is only allowed to displace in z and x directions, and h) the rigid merozoite is displaced in the negative y-direction.

The tight junction has the shortest element length relative to the erythrocyte membrane and is typically used to determine stable time increments for the entire model. For the erythrocyte membrane, represented using shell elements with a thickness of 0.01 µm (Hochmuth et al. 1973), a mesh density study was performed to determine the mesh density that provides a converged solution. SPH was applied to the erythrocyte cytoplasmic domain; hence, no mesh density study was performed for the cytoplasm. However, to accurately simulate the cytoplasm using the SPH functionality, Abaqus requires more than 10,000 particles. Hence, 18,000 particles were generated using the particle conversion functionality.

The erythrocyte membrane was meshed with 19,942 three-node triangular shell elements with reduced time integration (S3R). The reduced-time integration algorithm provided more accurate results while reducing the running time. Conventional shell elements were preferred because, unlike continuum shell elements, which only have displacement degrees of freedom, conventional shell elements have both displacement and rotational degrees of freedom. For the erythrocyte membrane to wrap around the merozoite effectively, each erythrocyte membrane node must have both displacement and rotation degrees of freedom. Erythrocytes undergo extreme deformations as they circulate through narrow capillaries in the human body. To accurately predict erythrocyte deformation, displacement and rotational degrees of freedom must be allowed in each element. Additionally, shell elements were preferred over 3D solid elements for the erythrocyte membrane because they enable the modelling of thin features with fewer elements, thereby reducing computational time. Shell elements are also easier to mesh and less prone to negative Jacobian errors, which can result from negative element volume or distorted elements that might occur when using extremely thin solid features.

The erythrocyte cytoplasm was meshed using 841 eight-node linear brick elements with reduced integration and hourglass control (C3D8R). The mesh provides the initial spatial particle discretisation required for the SPH scheme. Seven hundred three-node 3D rigid triangular elements (R3D3) were used for the rigid merozoite, and 5,535 ten-node modified quadratic tetrahedron elements (C3D10M) were used for the annulus structure of the tight junction.

### 2.4. Boundary conditions

Several assumptions and boundary conditions were considered:

- The erythrocyte was assumed to be suspended in an Euclidean space with an initial velocity of approximately zero for each node. Initial velocity fields were pre-defined at each node of the erythrocyte membrane (Figure 3 e).
- A constant external pressure load of 16 kPa, equal to systolic blood pressure, was applied on the external surface of the erythrocyte membrane (Figure 3 f).
- The merozoite was displaced by 2 µm along its longitudinal axis. The displacement was applied using a ramp function from 0.1 s to 1.1 s (Figure 3 g).
- Each node of the tight junction freely deformed in the x and z directions (Figure 3 h).

### 2.5. Contact interactions

Two algorithms were used to model contact interactions between structures involved in the invasion process. The general contact algorithm was used to define contact interactions between the erythrocyte membrane, the erythrocyte cytoplasm and the tight junction. The contact pair algorithm was used to define contact between the merozoite and the outer surface of the erythrocyte membrane. The contact pair algorithm in Abaqus Explicit includes the contact surface weighting (balanced or pure master-slave) and the sliding formulation (finite, small, or infinitesimal). The contact-pair algorithm with pure master-slave weighting was used to model contact between the merozoite surface and the region of entry (ROE) on the erythrocyte membrane. In the pure master-slave scheme, the interacting surfaces can penetrate each other, leading to numerical instabilities. To avoid numerical errors due to penetration, the mesh density of the slave surface must be greater than that of the master surface. A mesh density study was conducted for both surfaces to determine the optimal mesh sizes for the master and slave surfaces.

### 2.6. Model validation

#### 2.6.1. Validation of the erythrocyte finite element model with simulation of a healthy erythrocyte in an optical tweezer/trap

The mechanical response of the erythrocyte predicted by the developed model was compared to data from an optical trap experiment during which a force of 193 pN was applied to stretch the erythrocyte (Mills et al. 2004, Fig. 7). The stretching force was applied by trapping with a laser beam one of the two silica beads attached to the erythrocyte while the second silica bead was fixed to a glass slide while the other is (Song et al. 2017). Cell stretching is performed by moving the trapped microbead. The deformation was determined from images of undeformed and stretched erythrocytes.

During the finite element simulation of the optical trap experiment, an intact erythrocyte model (i.e. without membrane damage and with β₁ = β₂ = 0) was subjected to an axial tensile force of F = 200 pN. The force was applied to the FE nodes within a circular region of 1 µm in diameter on one side of the membrane, representing the adhesion interface between the membrane and the silica bead. Rigid body motion of the erythrocyte was prevented by constraining the FE nodes of a circular region on the axially opposite membrane side, simulating the adhesion to the second silica bead. These loading and boundary conditions produced deformation of the erythrocyte in the axial and transverse directions.

#### 2.6.2. Validation of the merozoite invasion finite element model

Recently, Geoghegan et al. (2021) used lattice light-sheet microscopy (LLSM) with high spatiotemporal resolution to analyse the merozoite invasion into the erythrocyte by segmenting and tracking the formation of parasitophorous vacuole membrane (PVM). The authors determined the portion of the erythrocyte membrane surface area that wraps the merozoite, and that does not. The study documents a decrease in erythrocyte membrane surface area, as a portion of the area wraps the merozoite (Geoghegan et al., 2021, Fig. 2c). The data obtained from this study were used to validate the developed invasion model. *In vivo*, erythrocytes are subjected to physiological blood pressure. Hence, blood pressure must be applied to the erythrocyte model surface to obtain realistic simulation results with the invasion model. However, no physiological pressure was used in the Geoghegan et al. (2021) experiment. As such, surface area data for the erythrocyte obtained from the invasion model in which no surface pressure was applied were compared with erythrocyte areal data from the Geoghegan et al. (2021) experiment to validate the invasion model.

### 2.7. Finite element analysis and case studies

#### 2.7.1. Generalised explicit finite element analysis in Abaqus

A generalised Abaqus Explicit dynamic analysis procedure was used to simulate the deformation of an erythrocyte during merozoite invasion. This procedure involves numerically solving the momentum equilibrium equation using an explicit central difference time integration rule described by Eqns. (40) and (41). The momentum equilibrium is:

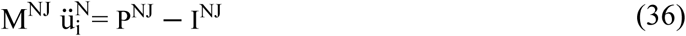

where M^NJ^ is the mass matrix, P^NJ^ is the applied load vector, I^NJ^ is the internal force vector, and ü^N^ denotes the spatial degrees of freedom. M^NJ^, P^NJ^ and I^NJ^ are defined as:

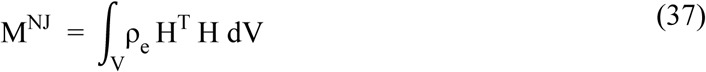

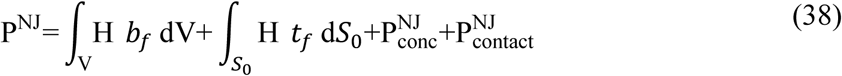

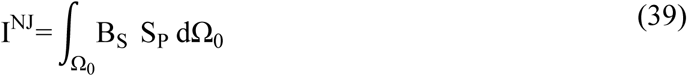

where ρ_e_ is the element density, Н is the shape function of nodes, b_f_ is the body force per unit mass, t_f_ is the pressure vector component, 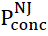 is the concentrated nodal load, and 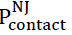 is the nodal contact force generated automatically by Abaqus contact algorithms, B_s_ is the strain-displacement matrix, and S_P_ is the second Piola–Kirchhoff stress. M^NJ^ is diagonalised to form a lumped mass matrix, thereby reducing the computational complexity of the central difference time integration algorithm (Dassault Systèmes Simulia Corp 2015).

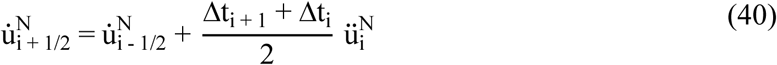

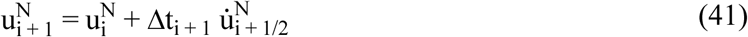

The dynamic explicit analysis with mass scaling and geometrically nonlinear analysis was used. Mass scaling was only applied to the erythrocyte membrane and the tight junction to obtain a quasi-static response. The quasi-static analysis was achieved by simulating the invasion process in the shortest time, i.e. 1.1 s, while keeping inertial forces relatively low. The semi-automatic mass scaling was used throughout the step to scale mass elements periodically and effectively reduce the wave speed. The effectiveness of the mass-scaling algorithm in ensuring a quasi-static solution was determined by confirming that the total kinetic energy of the erythrocyte model was much smaller than its internal or strain energy. Kinetic energy reflects the effects of inertia on the erythrocyte model’s global response, while internal energy reflects its static effects. Time incrementation was achieved automatically by using built-in functionality. The adaptive, global estimation algorithm was applied to determine the maximum frequency of the entire model using the current dilatational wave speed.

#### 2.7.2. Case Study 1: Impact of the erythrocyte morphology on the merozoite invasiveness

Using the developed finite element model, the impact of erythrocyte morphology and merozoite-induced damage on merozoite invasiveness was assessed. The invasiveness of the merozoite in the convex region (Figure 4 a) and the concave region (Figure 4 b) was assessed by comparing the total invasion energies associated with the merozoite’s invasion of the convex and concave regions. To assess the impact of sphericity, i.e., surface area to volume ratio, the total invasion energies were compared for the entry of the merozoite into a normal discoid-shaped erythrocyte and into a spherical erythrocyte with the same volume as the normal erythrocyte (Figure 4 a). To date, it is unknown whether the spherical shape impacts the invasiveness of the merozoite. The morphometric parameters used to define the two erythrocyte morphologies, i.e., discoid and spherical morphologies, are given in Table 1. The developed spherical model of the erythrocyte represents a 27% reduction of a healthy erythrocyte’s surface-to-volume ratio S/V and total surface area. With the two models of the erythrocyte, i.e., spherical and discoid models, the impact of morphological variations on the invasiveness of the malaria parasite was investigated.

**Figure 4:**
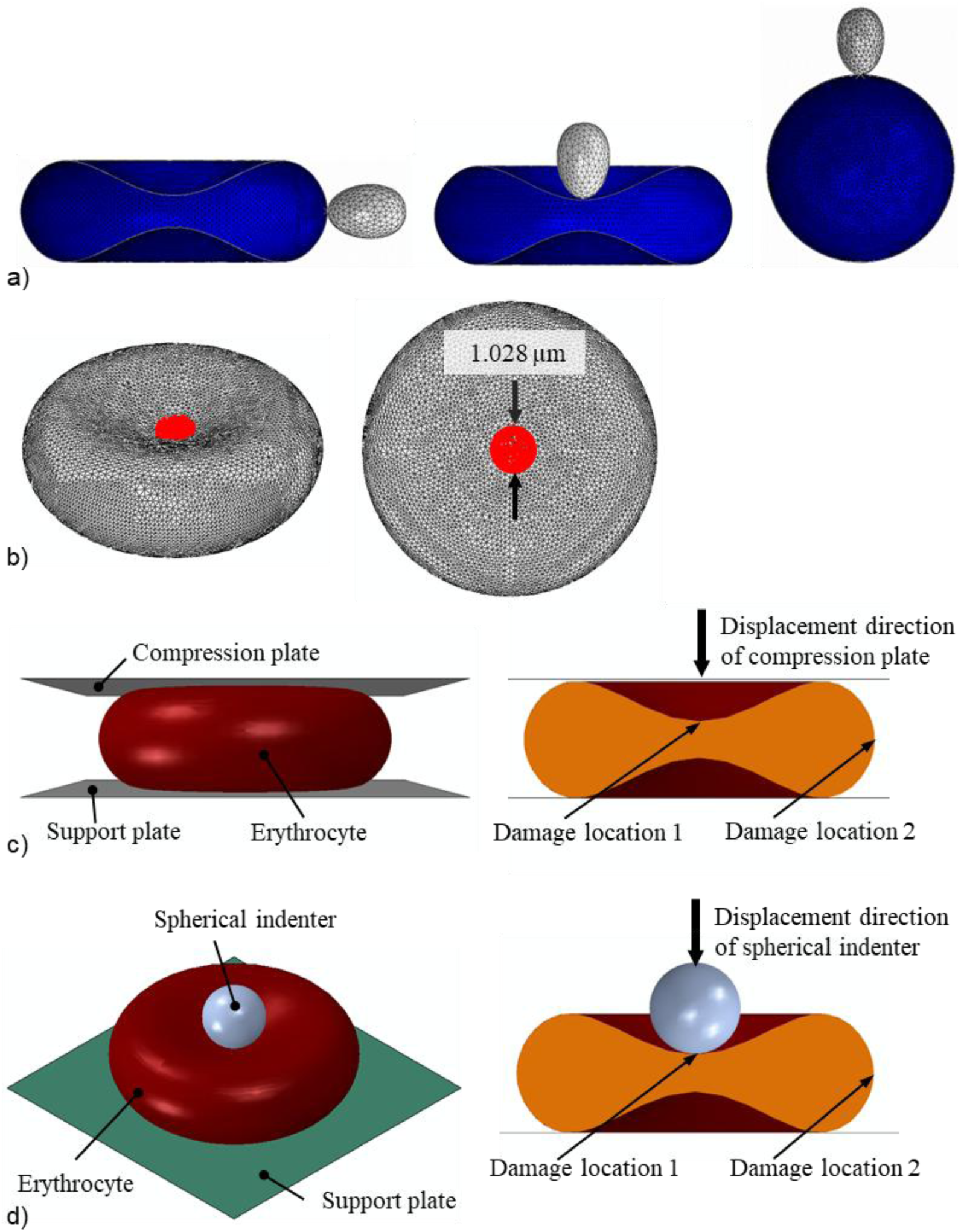
a) Case study 1: Entry point of the merozoite (grey) into a human erythrocyte (blue) in a convex (left) and concave region (middle), and into a human spherocyte (blue) (right). b) Case study 2: Location and dimension of damage (red) induced in erythrocyte membrane. e) Case study 3: Initial configuration of the compression test simulation (left) and locations of induced membrane damage in the concave and convex regions of the erythrocyte. d) Case study 4: Initial configuration of the nanoindentation simulation and locations of induced membrane damage in the concave and convex regions of the erythrocyte.

**Table 1:**
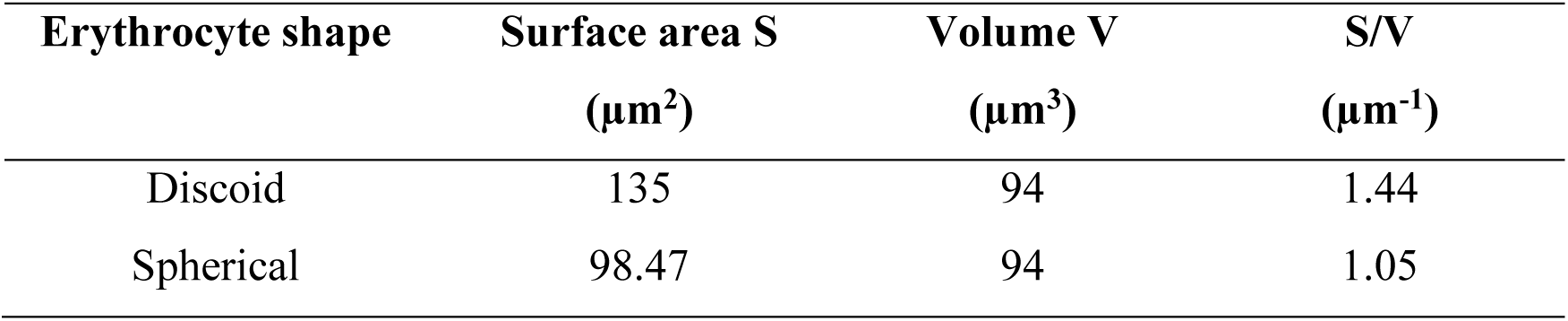
Erythrocyte parameters for case study 1.

| **Erythrocyte shape** | **Surface area S**<br>**($\mu\text{m}^2$)** | **Volume V**<br>**($\mu\text{m}^3$)** | **S/V**<br>**($\mu\text{m}^{-1}$)** |
| --- | --- | --- | --- |
| Discoid | 135 | 94 | 1.44 |
| Spherical | 98.47 | 94 | 1.05 |

#### 2.7.3. Case Study 2: Impact of phosphorylation-induced damage in the erythrocyte membrane on the merozoite invasiveness

Using the developed erythrocyte membrane damage model, the impact of the damage-induction period on merozoite invasiveness was assessed. Damage induction limited to the early invasion, i.e., τ = 0.1 s, was compared to damage induction throughout the entire invasion duration of τ = 1.1 s, where the amount of induced damage, *e*^β1τ^, was the same for both induction periods, *e*^β1τ^|_τ=0.1*s*_ = *e*^β1τ^|_τ=1.1*s*_. Damage was induced in a central circular region of the erythrocyte membrane with a diameter of 1.028 µm (Figure 4 b). For each damage induction period, the invasiveness of the merozoite was evaluated using the indentation force generated by the merozoite. The corresponding damage parameters used were β_1_ = 5.4 and 11 for τ = 0.1 s, and β1 = 0.49 and 1 for τ = 1.1 s.

#### 2.7.4. Case Study 3: Erythrocyte compression

The compression simulations with intact erythrocytes and erythrocytes with local membrane damage investigated the impact of local membrane damage on erythrocyte global mechanical and structural properties (Figure 4c). First, an intact erythrocyte model was compressed to extract the compression data. Secondly, an erythrocyte with a locally damaged erythrocyte membrane was compressed to extract two compression data sets, one obtained by inducing damage in the concave region, i.e., damage location 1 and the other in the convex region of the erythrocyte membrane, i.e., damage location 2 (Figure 4 c). The diameter of the damage region was 1.028 µm. The compression plate (10 × 10 µm) was displaced by 1.3 µm for a period of 1.1 s, while the support plate (10 × 10 µm) was fixed to facilitate compression of the erythrocyte. The compression and support plates were modelled using 5,000 linear triangular rigid shell elements (R3D3). The contact pair algorithm was used to define contact between the compression plate-erythrocyte interface and the erythrocyte-support plate interface, where surface-to-surface contact formulation with the kinematic contact method was defined. The normal and the tangential behaviour of the interfaces mentioned above were defined for this type of interaction. An isotropic tangential interaction with a negligible friction coefficient was used to describe the tangential behaviour at the interfaces. Hard contact formulation, which allows separation of the interfaces mentioned above, was used to define the normal behaviour of the interfaces. The force-compression data obtained from indenting a locally damaged and intact erythrocyte were compared to assess the sensitivity of the global indentation for local erythrocyte membrane damage. Sufficient sensitivity may indicate the potential of compression tests for further development and implementation to study merozoite-induced damage in a physical experiment. The impact of surface pressure on the mechanical response of the locally damaged erythrocyte membrane was also investigated.

#### 2.7.5. Case Study 4: Erythrocyte nanoindentation

The nanoindentation simulation was conducted using a similar approach to that of the compression test simulation in the previous section. The nanoindentation simulation involved an intact erythrocyte and an erythrocyte with a locally damaged membrane to investigate the impact of local membrane damage (Figure 4d). The spherical indenter with a diameter of 2 µm, representing 25.6% of erythrocyte diameter, was displaced by 1.31 µm, representing 64.7% of the maximum erythrocyte thickness. The support plate (10 × 10 µm) was fixed to facilitate indentation of the erythrocyte. The spherical indenter was modelled using 854 linear rigid triangular shell elements (R3D3), while the support plate was modelled using 5,000 linear rigid triangular shell elements (R3D3). The contact pair algorithm was used to define contact between the spherical indenter-erythrocyte interface and the erythrocyte-support plate interface, where the surface-to-surface contact formulation with the kinematic contact method was defined. Both normal and tangential behaviour of the interfaces mentioned above were defined for this type of interaction. An isotropic tangential interaction with a negligible friction coefficient was used to describe the tangential behaviour at the interfaces. Hard contact formulation that allows separation of the interfaces was used to define the interfaces’ normal behaviour. The force-indentation data obtained from the locally damaged and intact erythrocytes were compared to assess the sensitivity of the nanoindentation to local erythrocyte membrane damage. A sizeable difference between the force-indentation curves of the undamaged and damaged erythrocyte suggests that the nanoindentation test is sensitive to erythrocyte membrane damage.

Finite element simulations were conducted on the University of Cape Town High Performance Computing (HPC) facility using Abaqus on Windows-based multi-core compute nodes. CPU and memory resources were allocated through the SLURM scheduler.

## 3. Results

### 3.1. Erythrocyte membrane damage model

The true stresses defined by the Abaqus built-in Mooney Rivlin model and the erythrocyte membrane damage model with the parameter values provided in Table 2 agreed well for the intact membrane, i.e. for β_1_ = 0, and for true strain between 0 and 1.1 (Figure 5 a). This agreement indicates that the developed VUMAT subroutine was accurately implemented.

**Figure 5:**
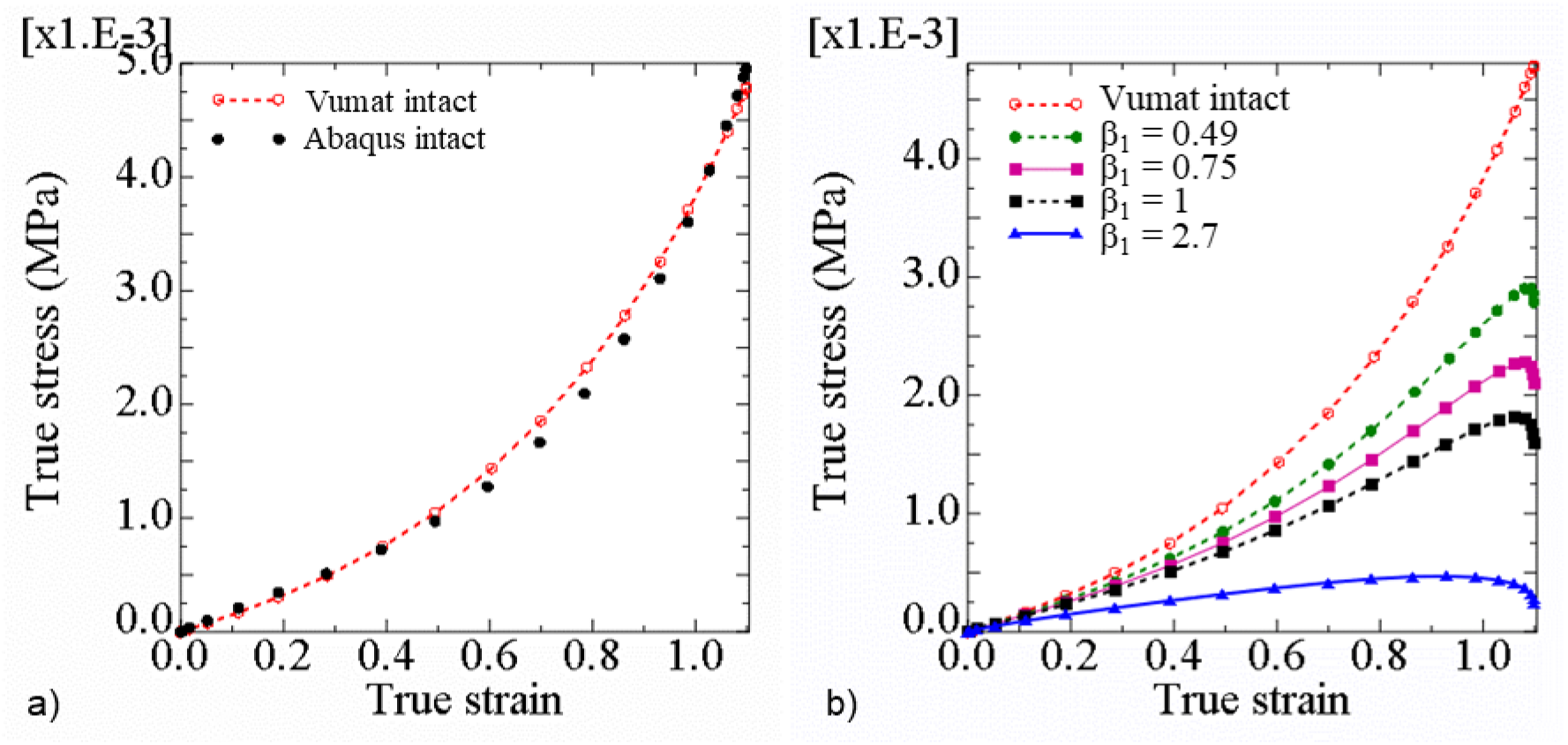
(a) True stress versus true strain in a shell element determined with the Abaqus built-in Mooney Rivlin model (‘Abaqus intact’) and the erythrocyte membrane damage model (‘Vumat intact’) for β_1_ = 0, and (b) true stress versus true strain determined with the erythrocyte membrane damage model showing the stress decrease with initiation and increase of erythrocyte membrane damage from β_1_ = 0 to 2.7.

**Table 2:** Material parameter values to represent an intact erythrocyte membrane with the erythrocyte membrane damage model.

| $D_1$ (mm<sup>2</sup>/N) | $C_{10}$ (μN/mm<sup>2</sup>) | $C_{01}$ (μN/mm<sup>2</sup>) | $\beta_1$ |
| --- | --- | --- | --- |
| 12 | 152 | 15.2 | 0 |

When the damage parameter was changed from the intact state with β_1_ = 0 to the damaged state with β_1_ = 2.7, the in-plane true stress decreased for any given true strain (Figure 5 b). For a true strain of 1.1, the in-plane true stress decreased from 0.0048 MPa for β_1_ = 0 to 0.0028 MPa for β_1_ = 0.49, 0.0016 MPa for β_1_ = 1, and 0.00025 MPa for β_1_ = 2.7.

### 3.2. Validation of the erythrocyte and invasion models

The developed erythrocyte model was validated by comparing optical tweezer experimental data (Mills et al. 2004) with numerical results from simulations of an optical tweezer experiment. When a force of 200 pN was applied diametrically, the dimension of the erythrocyte increased in the direction of the applied load, i.e. axial dimension (Figure 6 a) and decreased in the direction normal to the applied force, i.e. traverse dimension.

**Figure 6:**
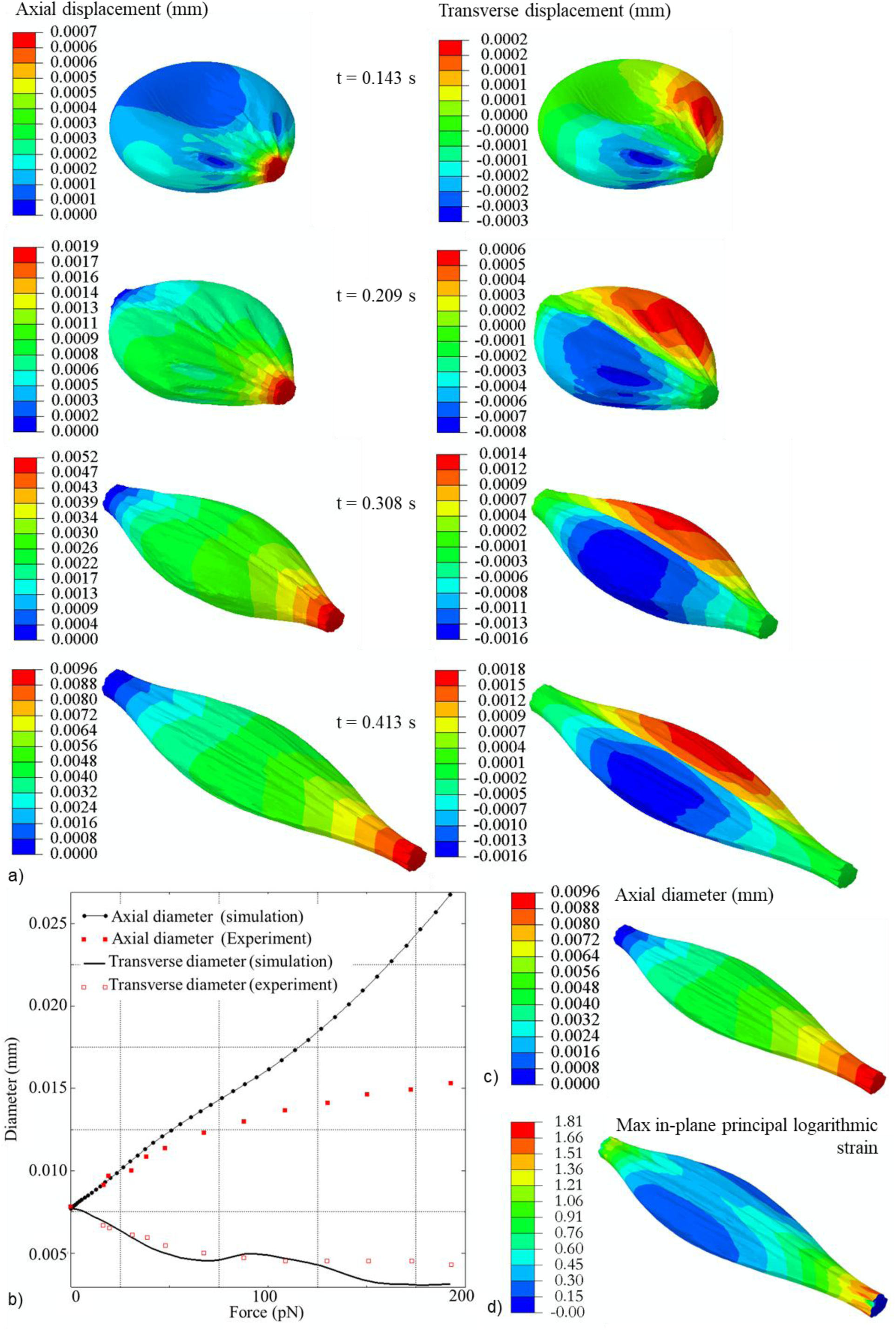
a) The contour plots of axial and transverse displacement of the erythrocyte predicted with the FEM model at 0.0143 s, 0.209 s, 0.308 s, and 0.413 s of the optical tweezer simulation. b) Optical tweezer simulation data (axial and transverse erythrocyte finite element model diameters) for an erythrocyte without membrane damage (i.e. β_1_ = 0 and β_2_ = 0) fitted with experimental data for the axial and transverse diameter of a human erythrocyte from Mills et al. (2004). Contour plots of axial diameter (c) and maximum principal logarithmic strain (d) of the erythrocyte predicted with the finite element model of an optical tweezer test.

**Figure 7:**
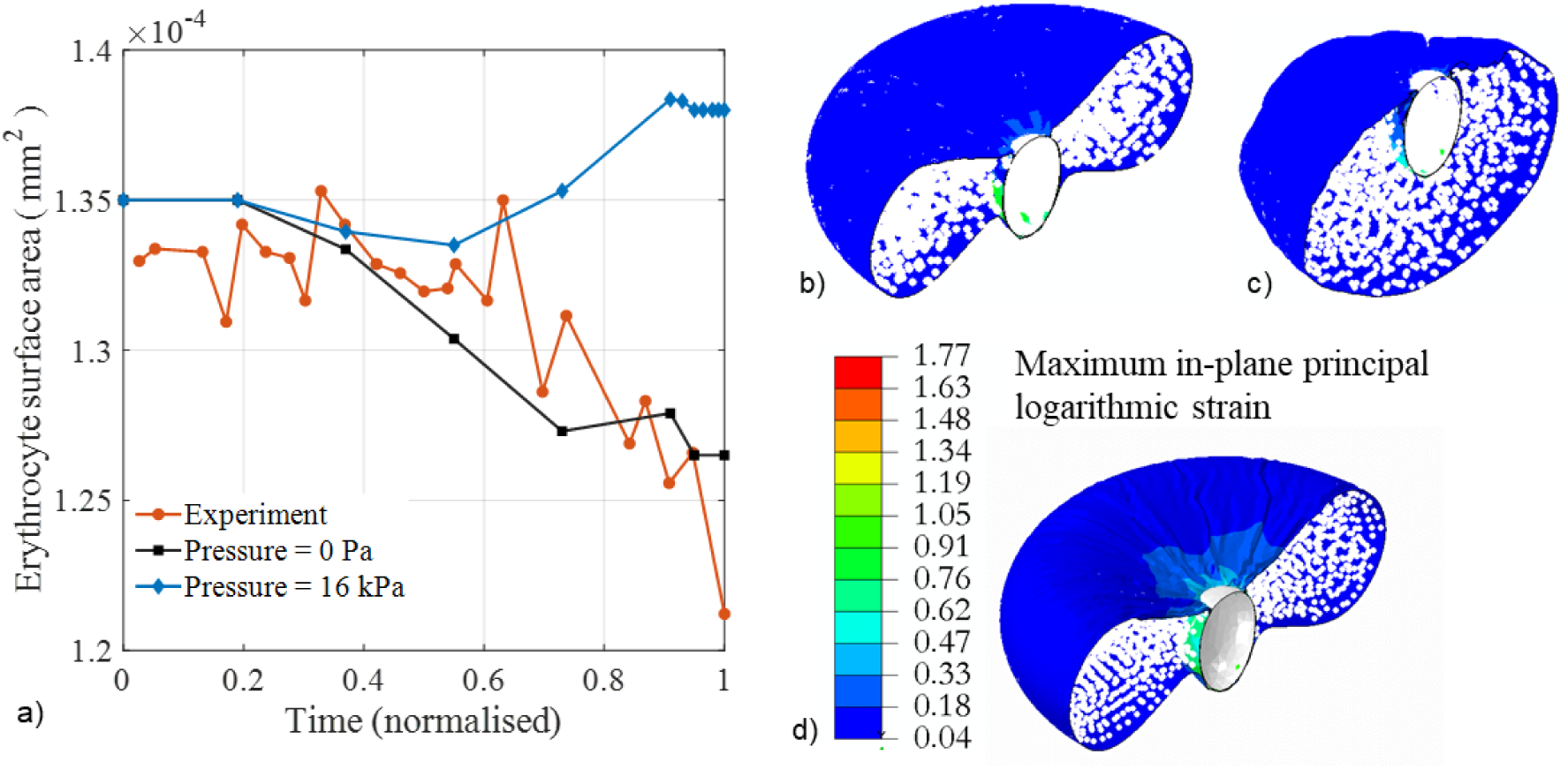
a) Variation of erythrocyte surface area during merozoite invasion predicted with the proposed invasion model with (blue diamonds) and without blood pressure (black squares), and experimental data from Geoghegan et al. (2021) (red circles) obtained without blood pressure. Erythrocyte membrane damage was modelled with β_1_ = 11 induced during early invasion with τ = 0 to 0.1 s. c and d) Deformation of an erythrocyte at 100% indentation depth normalised over the length of the merozoite with (b) and without (c) blood pressure applied on the outer surface of the erythrocyte. d) Contour plot of the maximum principal logarithmic strain in the erythrocyte membrane with blood pressure at 100% normalised indentation depth.

The increase in displacement of the erythrocyte finite element model in the axial direction leads to an increase in the axial diameter of the erythrocyte. In contrast, the increase in the transverse displacements of the erythrocyte finite element model leads to the reduction of the transverse diameter of the erythrocyte (Figure 6 a).

The numerical data show that the transverse diameter of the erythrocyte model fits well with the experimental data. However, the axial diameter of the erythrocyte model only fits well with experimental data when it is less than 0.0096 mm (Figure 6 b and c). For the erythrocyte model, the axial diameter of 0.0096 mm corresponds to the maximum principal erythrocyte membrane logarithmic strain of 1.81 (Figure 6 d). Beyond this point, the model fits experimental data in the axial direction with limited accuracy.

The invasion model was validated by comparing the erythrocyte surface area with experimental erythrocyte areal deformation data obtained by tracking and segmenting the erythrocyte membrane during the invasion process (Geogheg*an et a*l. 2021). The erythrocyte surface area numerically predicted for the case without blood pressure agrees well with the experimental data from Geoghegan et al. (2021), which were also obtained without blood pressure on the erythrocyte (Figure 7a). The maximum error between the experimental areal data (Geoghegan et al. 2021) and the numerically predicted data was 5.2%. Without blood pressure, the model predicts a decrease in erythrocyte surface area consistent with experimental data from Geoghegan et al. (2021). In contrast, when a blood pressure of 16 kPa is applied, the erythrocyte’s surface area increases during the late stages of the invasion process (Figure 7 a). The deformation of the erythrocyte model with a blood pressure of 16 kPa is entirely different from that without blood pressure (Figure 7 b and c). The largest maximum principal logarithmic strain in the erythrocyte membrane during the merozoite entry is 1.77 (Figure 7 d), which is less than the maximum principal logarithmic strain accuracy threshold of 1.81 established for the erythrocyte model.

### 3.3. Impact of erythrocyte morphology on merozoite invasiveness

The impact of morphological variations of the erythrocyte on the invasiveness of the merozoite is determined by the maximum amount of energy required for successful invagination of the merozoite (case study 1). When the merozoite invades a convex region of the erythrocyte, the maximum strain energy predicted by the invasion model is 0.38 × 10^-15^ J, whereas when it invades the concave region of the erythrocyte, the maximum strain energy of 0.238 × 10^-15^ J is predicted (Figure 8 a). For the invasion of a spherocyte, a maximum strain energy of 0.545 ×10^-15^ J is predicted, which is 43% and 129% larger than the energy required for the invagination of a merozoite in the convex and concave region of the erythrocyte, respectively.

**Figure 8:**
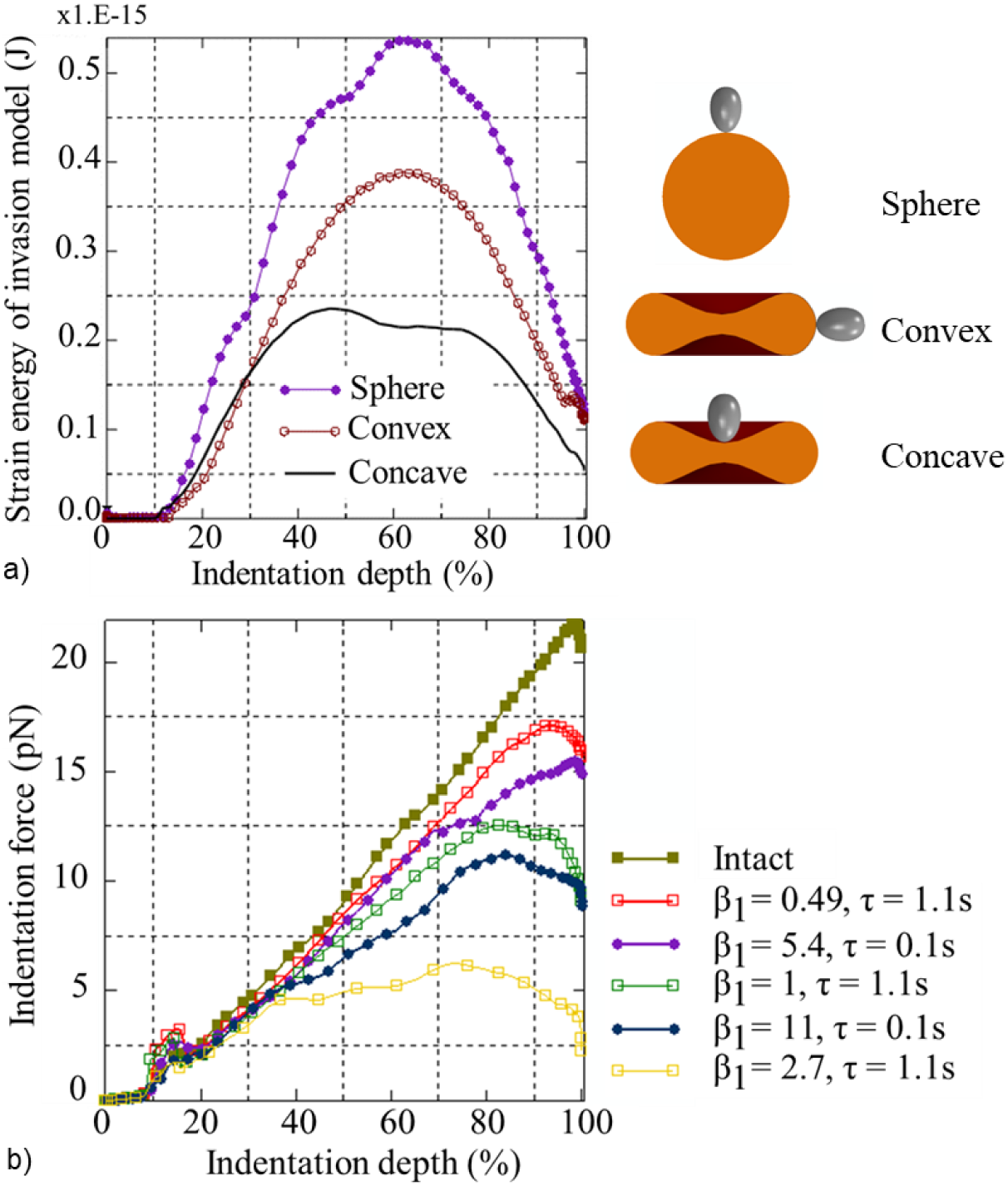
a) Required strain energy versus indentation depth predicted with the invasion model for invagination of a merozoite in the concave and convex region of the erythrocyte and the spherocyte. b) Indentation force versus indentation depth for varying degrees and timing of erythrocyte membrane damage.

### 3.4. Impact of phosphorylation-induced damage in the erythrocyte membrane on the merozoite invasiveness

The impact of malaria-induced erythrocyte membrane damage on the invasiveness of the merozoite (case study 2) is determined by the variation of the maximum indentation force required by the merozoite to invade the erythrocyte for increasing membrane damage represented by increasing β_1_ from 0 to 2.7 (Figure 8 b).

For erythrocyte membrane damage, *e*^β1τ^, limited to the early invasion stage (i.e. for τ = 0.1 s of the total simulation time of τ = 1.1 s), the maximum indentation forces are lower than for an equal amount of erythrocyte membrane damage induced for the entire simulation time. For example, the damage induced for β_1_ = 0.49 with τ = 0.1 s is the same as for β_1_ = 5.4 with τ = 1.1 s. Similarly, equal damage is obtained for β_1_ = 1 with τ = 1.1 s and β_1_ = 11 with τ = 0.1 s. The maximum indentation force is larger for an intact erythrocyte membrane (22.4 pN) than for an erythrocyte with membrane damage of β_1_ = 0.49 and τ = 1.1 s (17 pN). When the same damage amount is induced in the erythrocyte membrane during early invagination for τ = 0.1 s (with β_1_ = 5.4), the maximum indentation force is yet lower (15 pN). Similarly, the maximum indentation force of 11 pN for β_1_ = 11 with τ = 0.1 s is lower than the force of 12.5 pN for β_1_ = 1 with τ = 1.1 s.

### 3.5. Impact of local erythrocyte membrane damage on the global mechanical responses of the erythrocyte

For compression of the entire erythrocyte (case study 3), the force-displacement curves do not differ between the normal erythrocyte and that with damage at location 1 and 2, respectively, with β_1_ = 32 for τ = 0.1 s (Figure 9 a). This result shows that global compression is insensitive to local damage to the erythrocyte membrane caused by phosphorylation.

**Figure 9:**
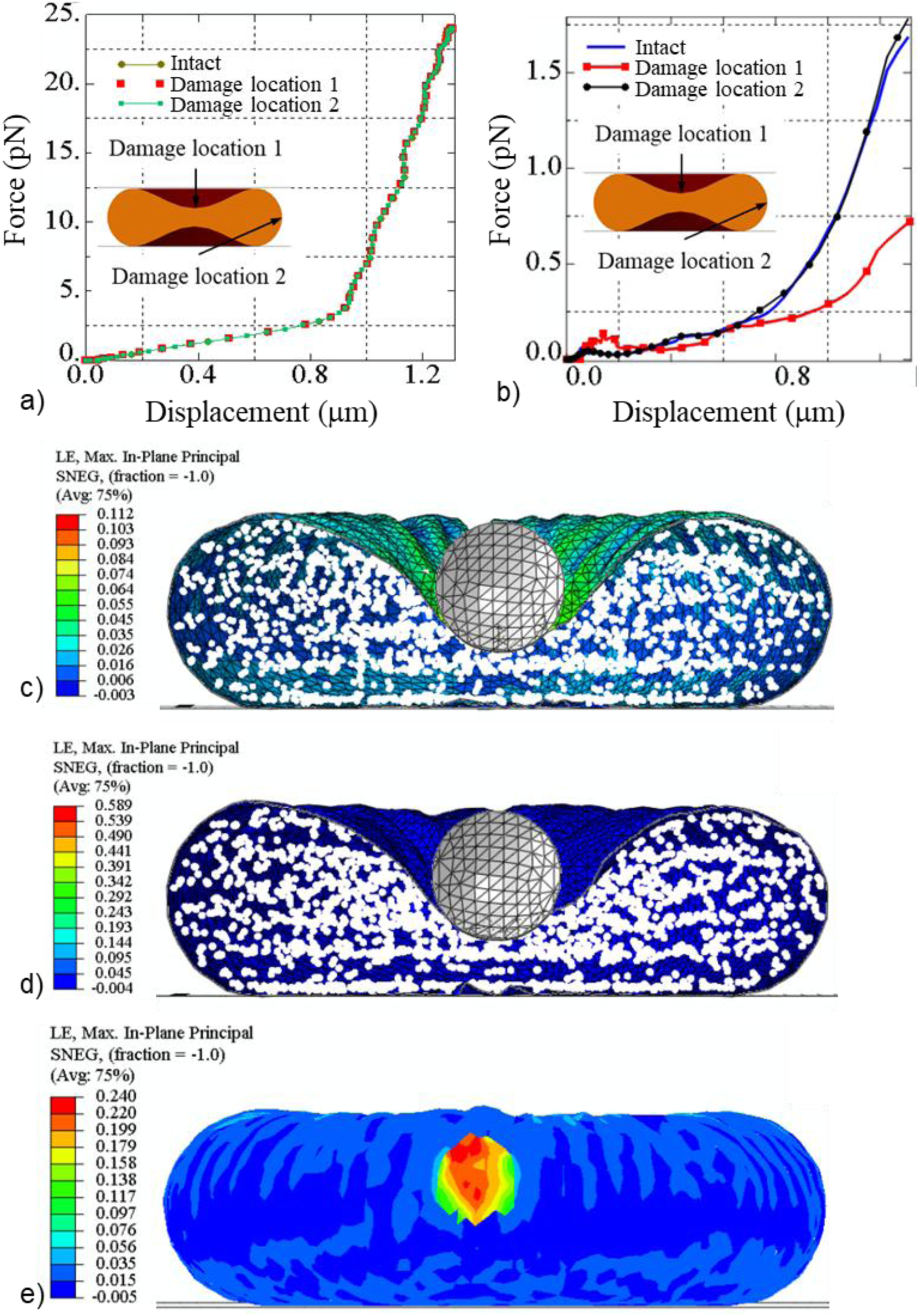
Predicted force versus displacement for compression (a) and nanoindentation (b) of an intact erythrocyte and an erythrocyte damaged at damage locations 1 and 2, respectively. Maximum principal logarithmic strain for the intact erythrocyte membrane (c). Maximum principal logarithmic strain for the erythrocyte membrane with damage (β_1_ = 32) induced for s at damage location 1 (d) and 2 (e) where A and A_s, max_ denotes the surface area and the areal ratio of the erythrocyte, respectively.

For local nanoindentation of an intact erythrocyte (case study 4), the maximum indentation force is 2.5-fold lower for damage induced at damage location 1 (i.e. the region of the indentation) (0.72 × 10^-12^ N) than at damage location 2 (1.78 ×10^-12^ N) (Figure 9 b). The latter is similar to the maximum indentation force for the intact erythrocyte membrane (1.68 ×10^-12^ N). An indentation displacement of 1.31 µm causes a maximum principal logarithmic strain in the erythrocyte membrane of 0.112 without blood pressure (Figure 9 c). For damage induced at damage locations 1 and 2, respectively, with β_1_ = 32 for 0.1 s, the maximum principal logarithmic strain is 0.589 (Figure 9 d) and 0.24 (Figure 9 e). The surface area of the intact and the damaged erythrocytes increases equally from 135 µm^2^ to 137 µm^2^, resulting in an areal strain of A_s,max_ = 0.014 (Figure 9 c to e).

## 4. Discussion

Although considerable progress has been made in understanding which factors determine merozoite invasiveness, most studies have not addressed the collective role of the biophysical characteristics of erythrocyte deformability in the invasion process. Cell shape, cytoplasmic viscosity, and membrane stability are the main determinants of erythrocyte deformability (Mohandas and Chasis 1993; Mohandas and Evans 1994). Experimental data on the mechanical properties of the human erythrocyte are widely available; however, it remains unknown how these physical properties influence the invasiveness of the merozoite. Hence, in the current study, an *in silico* approach was developed to investigate the role of erythrocyte morphology and merozoite-induced damage to the erythrocyte membrane, which are difficult to isolate experimentally, in the invasiveness of the merozoite. Importantly, this approach provides a mechanistic understanding that contributes to the rational identification of biomechanical targets for cell-mechanics-based antimalarial drug screening (Krishnan et al. 2016).

### 4.1. Erythrocyte membrane damage model

To date, experimental investigations of erythrocyte invasion have predominantly focused on the roles of parasite adhesins, signalling pathways, and the identity of binding receptors on the erythrocyte surface. Erythrocyte membrane damage mechanics associated with the invasion process have received limited attention (Koch et al. 2017).

The erythrocyte membrane damage model developed in the current study utilises the hyperelastic Mooney Rivlin constitutive law, complemented with an exponential damage function to account for membrane remodelling due to phosphorylation. Despite limited knowledge of merozoite-induced damage, the model allowed for the representation of varying levels of damage. One constraint of hyperelastic constitutive models is their instability when strain is inversely related to stress. One way to assess a model’s stability is using Drucker’s criterion. However, this method has challenges, as some hyperelastic models can be Drucker-unstable at both small and large strains under different loading conditions. Hence, standard practice is to use a model with a known stability threshold or validity range. For example, the Mooney Rivlin model has a known validity range for equi-biaxial logarithmic strain up to 138% (Marckmann and Verron 2006).

The stability threshold of the developed constitutive damage model was analytically determined using Drucker’s stability criteria. The decrease in the maximum stress indicates a reduction in stiffness when damage is induced (Figure 5b), indicating that the developed damage model accurately mimics the phosphorylation of the key erythrocyte membrane skeleton. The main limitation of the developed erythrocyte membrane damage model is the possible instability of the underlying Mooney Rivlin law at low strain. Furthermore, the second invariant of the left Cauchy-Green tensor of the Mooney-Rivlin law can make the developed model unstable under certain loading conditions. However, the decrease in the erythrocyte membrane stability threshold predicted by the developed model for increasing damage amounts corresponds with reports that the phosphorylation of key proteins in the erythrocyte skeleton leads to an unstable erythrocyte membrane.

### 4.2. Erythrocyte and invasion finite element models

The developed erythrocyte model is a two-component model of the erythrocyte membrane and cytoplasm. The diametric dimensions of the erythrocyte predicted by the developed model for a simulated optical-trap experiment were compared with experimental data from Mills et al. (2004) to validate the erythrocyte model. The predicted transverse erythrocyte diameters agree well with the experimental transverse diameters from Mills et al. (2004). However, the predicted axial diameter fits the experimental data only for axial diameters smaller than 0.0096 mm, corresponding to a maximum principal logarithmic strain of 1.81 in the erythrocyte membrane. Beyond this threshold, the developed erythrocyte model shows limited accuracy.

The invasion model was validated by comparing the predicted erythrocyte surface area during merozoite invasion with experimental data from Geoghegan et al. (2021). Since the erythrocyte was not exposed to blood pressure during the *in vitro* experiments, blood pressure was neglected in the invasion model for the validation simulation. The numerically predicted surface area excludes the region of the erythrocyte membrane that is in contact with the merozoite during wrapping. To allow direct comparison of the change in surface area during the invasion, the predicted data were normalised to τ = 1.1. In contrast, the experimental data was normalised to the time of 49 s required for complete internalisation of the merozoite in the erythrocyte (Geoghegan *et al*. 2021). The predicted and experimental erythrocyte surface areas agreed well (Figure 7a). The validated model allows prediction of the invasion process under *in vivo* conditions, e.g., blood pressure acting on the erythrocyte, which may be more challenging to achieve in *in vitro* experiments.

The validation of the invasion model also validates the developed erythrocyte membrane damage model, as the model predictions were in good agreement with the experimental data for β_1_ = 11 and τ = 0.1 s. Identifying the damage parameters enables the quantification of the correct invasion forces required for the merozoite to invade the erythrocyte. The invasion model indicated a maximum principal logarithmic strain in the erythrocyte membrane of 1.77. This value is below the erythrocyte model’s accuracy strain threshold of 1.81, indicating that the erythrocyte deformation during the invasion process is within the determined accuracy range of the erythrocyte model.

### 4.3. Impact of the erythrocyte morphology on the merozoite invasiveness

The implication of morphological variations of the erythrocyte on the invasiveness of the merozoite was assessed by comparing the invasion energetics for (i) a discoid erythrocyte and a spherocyte and (ii) for the concave and convex membrane region of the discoid erythrocyte.

The merozoite entry requires lower invasion energy for the discoid erythrocyte than the spherocyte. Reducing the surface area to volume ratio (S/V) increases the sphericity of the erythrocyte and leads to the formation of the spherocyte. The relatively low energy requirement indicates that the merozoite is more invasive when it invades a discoid-shaped erythrocyte than a spherocyte. An increase in sphericity corresponds to an increase in the energy required for the merozoite to invade the erythrocyte. The S/V of 1.44/m allows a healthy erythrocyte to undergo a large deformation of up to 230% of its original dimension. Reducing the healthy erythrocyte’s S/V by 14% forms a spherocyte with a surface area of 98.5 µm^2^ compared to the surface area of 135 µm^2^ of a healthy erythrocyte. The discoid erythrocyte shape provides an excess surface area of 36.5 µm^2^, i.e. 4.6-fold the surface area of 8.0 µm^2^ of a merozoite (Dasgupta et al. 2014), sufficient to facilitate the wrapping of the merozoite. The maximum strain energy predicted by the developed finite element invasion model matches the total indentation work predicted by our analytical model (Msosa et al. 2023). The maximum strain energy of 38.0 × 10^-17^ J and 23.8 × 10^-17^ J predicted with the finite element invasion model for invasion in the convex and concave erythrocyte membrane region, respectively, is larger than the total indentation work of E_i_ = 1.40 × 10^-17^ J predicted by the analytical model for an areal strain of A_s,max_ = 51%. The higher strain energy predicted by the finite element invasion model may be due to deformation of the erythrocyte cytoplasm, which is not accounted for in the analytical model.

Erythrocytes with membrane protein abnormalities, such as hereditary spherocytosis, are generally spherical and less deformable than normal discoid erythrocytes. However, it is unknown whether these alterations may present the merozoite with a less ideal condition for invasion of erythrocytes by merozoites. Spherocytes have been found to be less susceptible to invasion by merozoites. One reason for low susceptibility is genetic alterations in membrane proteins. However, the current study found that erythrocyte shape alteration to a spherical shape could be one of the contributing factors to the low susceptibility of spherocytes to merozoite infection.

### 4.4. Impact of phosphorylation-induced damage in the erythrocyte membrane on merozoite invasiveness

The impact of erythrocyte membrane damage on merozoite invasiveness was studied by inducing local erythrocyte membrane damage. The damage amount in the erythrocyte membrane model was regulated by varying damage parameter β_1_ between 0.49 and 2.7 such that β_1_ = 0.49 represented the minimum amount of damage and β_1_ = 2.7 represented the maximum amount of damage. The invasiveness of the merozoite was assessed by comparing the maximum indentation forces for each value of the damage parameter β_1_.

The indentation force decreases with an increase in the amount of damage in the erythrocyte membrane model (Figure 9). This demonstrates that the merozoite invasiveness increases with the extent of damage. Merozoite-induced erythrocyte membrane damage has received limited attention, and the stages of erythrocyte membrane remodelling or damage are unknown. It is also unknown whether damage is induced only at the early invasion stage or throughout the invasion process. To validate the developed invasion model, erythrocyte membrane damage was induced at the beginning of the invasion process, i.e. at τ = 0.1 s with β_1_ = 11. The results suggest that erythrocyte membrane damage occurs during an early stage of invasion. The merozoite requires a greater force when damage is induced progressively throughout the invasion, i.e., for τ = 1.1 s, than at the beginning of the invasion process with τ = 0.1 s. These results demonstrate that the merozoite is more invasive when damage is induced during the early invasion stage (τ = 0.1 s) than when it is induced progressively throughout the invasion process. Hence, regulating the timing at which the merozoite induces erythrocyte membrane damage could be a potential target for antimalarial compounds.

### 4.5. Impact of local erythrocyte membrane on the global mechanical responses of the erythrocyte

Compression simulations investigated the impact of local erythrocyte membrane damage on a global scale. The compression force does not differ for the intact and damaged erythrocyte, irrespective of the location of the phosphorylation damage (Figure 9 a). Hence, global compression of single erythrocytes cannot be used to detect erythrocyte membrane damage reliably. The simulations of nanoindentation of the erythrocyte in the central region indicate a discernible difference in the indentation force between an intact erythrocyte and an erythrocyte with membrane damage for damage in the central, concave region but not in the convex region of the discoid cell (Figure 9 e). This finding demonstrates that merozoite-induced local membrane damage can be detected by nanoindentation, depending on the damage location, and that further research is required.

## 5. Conclusions

In this study, a finite element invasion model was developed and used to (i) quantify the mechanics of a malaria merozoite’s invasion of an erythrocyte and (ii) investigate the impact of erythrocyte shape and membrane damage on the invasiveness of a malaria merozoite. The findings include the smallest force required for the malaria merozoite to invade a human erythrocyte, i.e., 11 pN successfully. The invasiveness of the merozoite decreases with increasing erythrocyte sphericity, which is associated with genetic disorders such as hereditary spherocytosis. An increase in phosphorylation-induced membrane damage in erythrocytes increases the invasiveness of the malaria merozoite, as might be expected. It was further found that the malaria merozoite is more invasive when erythrocyte membrane damage induced by phosphorylation is limited to the early invasion stage, rather than throughout the entire invasion stage. The findings on invasion mechanics can guide future experimental studies to assess the merozoite’s invasiveness. The results from the nanoindentation simulations indicate that nanoindentation is a suitable additional experimental technique for evaluating erythrocyte membrane damage in the context of invasion-blocking anti-malaria drugs. The computational models developed for human erythrocyte and merozoite invasion can be adapted to study other parasite invasion processes.

## Data availability

Software used and data supporting the results presented in this article are available on the University of Cape Town’s institutional data repository (ZivaHub) under https://doi.org/10.25375/uct.28263767 as Msosa C, Abdalrahman T, Franz T. Software code and data for “*In silico* analysis of the invasion mechanics and invasiveness of the plasmodium falciparum merozoite”, Cape Town, ZivaHub, 2025, DOI 10.25375/uct.28263767.

## CRediT author contributions

CM: Conceptualization, Data curation, Formal analysis, Funding acquisition, Investigation, Methodology, Project administration, Software, Validation, Visualization, Writing – Original Draft, and Writing – Review & Editing

TA: Conceptualization, Methodology, Project administration, Supervision, and Writing – Review & Editing

TF: Conceptualisation, Funding acquisition, Methodology, Project administration, Resources, Supervision, Validation, Visualization, and Writing – Review & Editing

